# *ybx1* acts upstream of *atoh1a* to promote the rapid regeneration of hair cells in zebrafish lateral-line neuromasts

**DOI:** 10.1101/2024.10.15.618534

**Authors:** Caleb C. Reagor, A. J. Hudspeth

## Abstract

Like the sensory organs of the human inner ear, the lateral-line neuromasts (NMs) of fish such as the zebrafish (*Danio rerio*) contain mechanosensory hair cells (HCs) that are surrounded by progenitors called supporting cells. Damaged NMs can quickly regenerate new HCs by expressing in the progenitors HC-specific genes such as *atoh1a*, the master regulator of HC fate. We used the supervised learning algorithm DELAY to infer regenerating NMs’ early gene-regulatory network (GRN) and identify adaptations that promote the rapid regeneration of lateral-line HCs in larval zebrafish. The central hub in the network, *Y-box binding protein 1* (*ybx1*), is highly expressed in HC progenitors and young HCs and can recognize DNA-binding motifs in cyprinids’ candidate regeneration-responsive promoter elements for *atoh1a*. We showed that NMs from *ybx1* mutant zebrafish larvae display consistent, regeneration-specific deficits in HC number and initiate both HC regeneration and *atoh1a* expression 20 % slower than in siblings. By demonstrating that *ybx1* promotes rapid HC regeneration through early *atoh1a* upregulation, the results support DELAY’s ability to identify key temporal regulators of gene expression.

## Introduction

Unlike humans, fish and amphibians can regenerate damaged receptors in the sensory organs of their inner ears as well as in their lateral-line organs for sensing “touch at a distance”^1–3^. During development, the lateral lines form when otic placode-derived cells migrate through the skin and deposit sensory units known as neuromasts (NMs) along the head, trunk, and tail^4–7^. Like inner-ear sensory epithelia, NMs contain two principal cell types: mechanosensory hair cells (HCs) for detecting directional displacement of water and HC progenitors called supporting cells (SCs), which surround the HCs^8–11^. Fish such as the zebrafish (*Danio rerio*) rely on sensory information from the lateral line to perform important behaviors such as rheotaxis and escape response^12,13^. However, due to their location near the surface of the skin, NMs often receive chemical and mechanical insults that damage HCs and leave larvae and adults vulnerable to predation^14^.

Because damaged NMs can regenerate HC progenitors within 3-5 hrs and mature HCs in less than 12 hrs, they are among the fastest-regenerating vertebrate sensory organs^15,16^. After injury, NMs recruit macrophages to clear debris and initiate proliferation of the SCs in their dorsal and ventral regions^17–20^. Depending on their expression of the transcription factor (TF) *atoh1a*— the master regulator of HC fate—the centrally located SCs may instead prepare to become unipotent HC progenitors^21–25^. The ability of NMs to quickly replace damaged HCs therefore depends on the maintenance of these progenitors and prompt induction of key HC-specific genes. The timing of *sox2* expression is also important because deletion of its upstream regeneration-responsive enhancer impedes HC regeneration^26^. Here we have sought additional gene-regulatory adaptations that foster the rapid regeneration of lateral-line HCs in the larval zebrafish. These mechanisms might present novel strategies for promoting HC regeneration in humans as well^27^.

## Results

### Supervised inference of the early GRN in regenerating NMs

We previously developed DELAY, a convolutional neural network for the identification of direct, causal interactions between TFs and their target genes from pseudotime-ordered single-cell gene expression trajectories^28^. DELAY employs transfer learning to reconstruct GRNs and identify key temporal hubs, such as *ESRRB* during the differentiation of Purkinje neurons^29^. We fine-tuned DELAY on single-cell RNA-sequencing (scRNA-seq) data of HCs and SCs immediately following NM injury, from zero to ten hours post ablation (hpa) of HCs^16^. After inferring the full HC- and SC-specific GRNs for 303 TFs expressed in either lineage, we identified the top twenty regulators for each gene and formed a network of the key TFs coordinating the early injury response and lineage specification (Figures 1A, 1B). The central hub, *Y-box binding protein 1* (*ybx1*), is a regulator of 255 TFs in the network.

**Figure 1:**
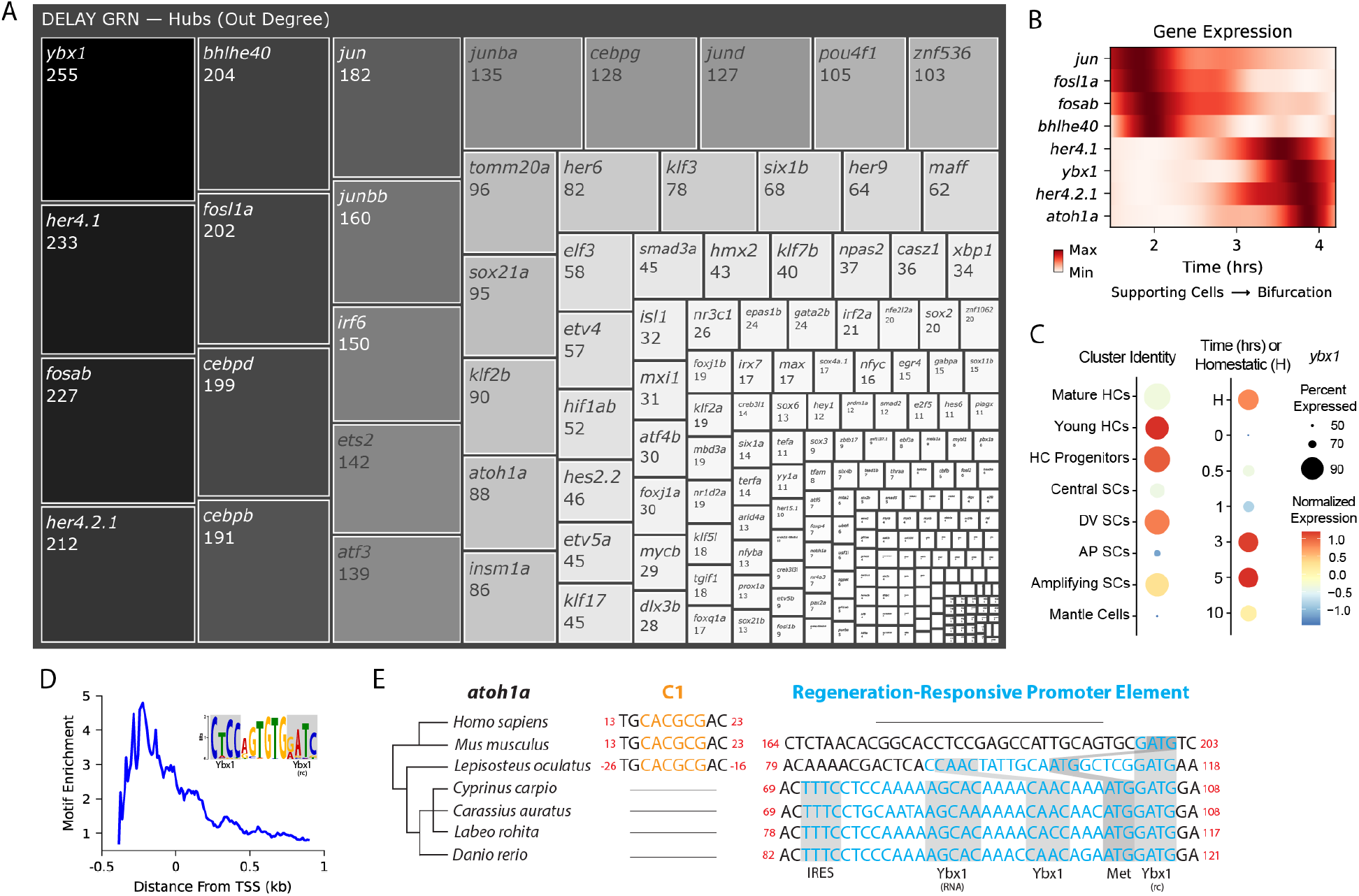
Centrality of *ybx1* during early HC regeneration. (A) A supervised GRN reconstruction with DELAY, arranged by outdegree centrality, identifies early gene-regulatory hubs for regenerating HCs and SCs in larval zebrafish NMs. (B) Generalized additive models depict the transition between injury-responsive gene expression (*jun, fosl1a, fosab*) and subsequent lineage specification (*her4*.*1, her4*.*2*.*1, atoh1a*) at 2-4 hpa. (C) A dot plot shows that *ybx1* mRNA is expressed across all cell types and times but is relatively enriched across HC progenitors and young HCs at 3-5 hpa. The analysis in A-C is based on data from ^16^. (D) *De novo* discovery of enriched motifs near targets’ transcription start sites (TSSs) suggests that Ybx1 binds to both forward and reverse DNA strands to regulate transcription. (E) Alignment of *atoh1a* reveals a candidate regeneration-responsive promoter element with two Ybx1 DNA-binding sites, a 5’ UTR element with an internal ribosome entry site (IRES) and Ybx1 RNA-binding site, and the loss of a repressive C1 site. rc, reverse complement.

Although no HC-specific roles are known, Ybx1 is a broadly expressed DNA- and RNA-binding protein that regulates transcription, translation, and cellular stress responses^30–32^. scRNA-seq detected *ybx1* mRNA in at least 50 % of all NM cells, and especially in HC progenitors and young HCs (Figure 1C)^16^. Because DELAY relies primarily on highly expressing cells, strong mRNA expression provided a robust signal to identify *ybx1*’s putative targets^28^. The neural network’s low false-positive rate also facilitated *de novo* discovery of enriched motifs across the enhancers of *ybx1*’s predicted targets. Proximally to its targets’ transcription start sites (TSSs), we discovered an enriched, bipartite motif that contained both forward and reverse-complement Ybx1 DNA-binding sites (Figure 1D). Ybx1 can bind to pyrimidine-rich sequences on single-stranded DNA (ssDNA)—often near other ssDNA-binding factors, or itself—to melt double-stranded DNA and regulate transcription of its targets^30,33,34^.

### A candidate regeneration-responsive promoter element for *atoh1a*

We noticed that one Ybx1 bipartite motif overlapped with the start codon of *atoh1a*, suggesting that *ybx1* operates upstream of HC specification. Alignment of the surrounding nucleotides revealed a candidate regeneration-responsive promoter element (cRRPE) in cyprinids—carps and minnows (Figure 1E). In addition to two Ybx1 DNA-binding sites, the cRRPE contains an element in the 5’ untranslated region (UTR) with a Ybx1 RNA-binding site—the so-called “dorsal localization element”—consisting of the sequence AGCAC followed by a short hairpin or stem-loop (CCA-N_6_-TGG), as well as an internal ribosome entry site (IRES)^35,36^. The IRE and RNA-binding sites suggest that Ybx1 can stimulate cap-independent translation of *atoh1a* mRNA while suppressing global, cap-dependent translation^34^.

The orthologous alignment of *atoh1a* across cyprinids including *Cyprinus carpio* (common carp), *Carassius auratus* (goldfish), and *Labeo rohita* (rohu) as well as other species such as *Lepisosteus oculatus* (spotted gar), *Mus musculus* (mouse), and *Homo sapiens* (human) suggests an origin for the cRRPE no later than the last common ancestor of the Neopterygii (carps, minnows, and gars), resulting from the emergence of a forward Ybx1 DNA-binding site near a putative reverse-complement site. Subsequent elaborations no later than the last common ancestor of the cyprinids resulted in the consolidation of spacing between DNA-binding sites, loss of the repressive C1 site for silencing by Hes/Hey-family Notch effectors, and inclusion of the 5’ UTR element^37^.

### *ybx1* promotes the rapid regeneration of HCs

To investigate the impact of *ybx1* on HC development and regeneration, we obtained *ybx1*^*sa34489*^ mutant zebrafish larvae that possessed an early stop in the ninth codon of *ybx1*. We inbred heterozygous adults to obtain wild-type, heterozygous, and homozygous mutant larvae, which we then treated with neomycin to ablate their lateral-line HCs (Figure 2A). At 5 days post-fertilization (dpf), we did not observe a difference in the number of HCs between the NMs of mutants and those of untreated siblings (Figure 2B). However, both the heterozygous and homozygous mutant NMs regenerated about 1.5 fewer HCs by two days post ablation (dpa) than in siblings (Figure 2B). *ybx1*^*sa34489*^ is therefore a dominant mutation that specifically disrupts HC regeneration in both heterozygous and homozygous mutant larvae.

**Figure 2:**
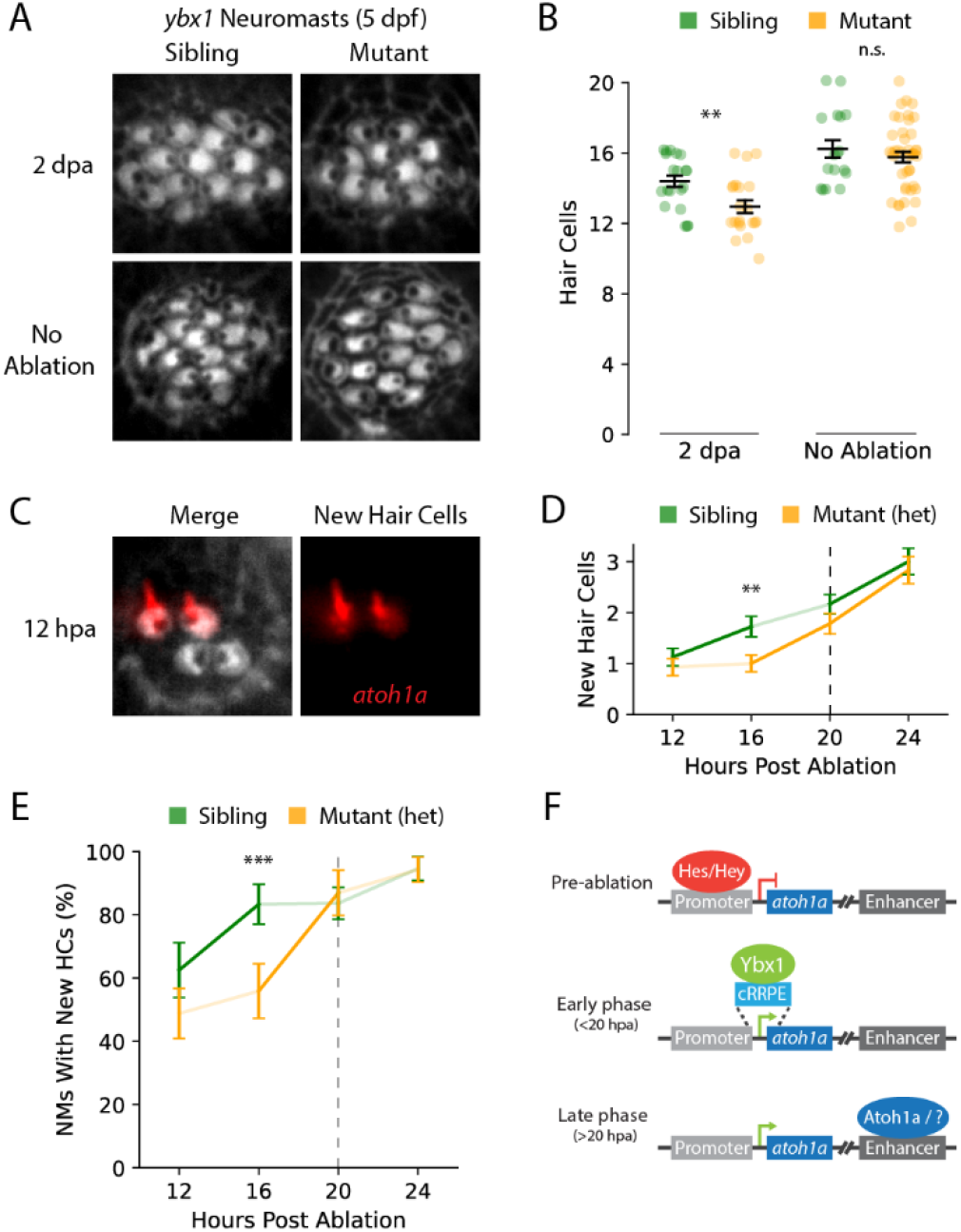
*ybx1* mutations delay *atoh1a* expression and HC regeneration. (A) Single-channel fluorescence images show phalloidin staining of HCs in post-ablation (top) and control (bottom) NMs, for either sibling (left) or heterozygous and homozygous *ybx1* mutant (right) larvae. (B) Average HC counts at 5 dpf are similar for control (right) sibling (n=17) and *ybx1* mutant (n=43) NMs, but neomycin-treated (left) mutants (n=21) regenerate about 1.5 fewer HCs than sibling (n=20) NMs by 2 dpa. (C) Two-channel fluorescence images show four HCs (left), including two newly regenerated *atoh1a*^*+*^ HCs (right) at 12 hpa. (D) Two lines show the average numbers of newly regenerated HCs in fixed, sibling NMs at 12 (n=32), 16 (n=36), 20 (n=55), and 24 hpa (n=37) and heterozygous mutants at 12 (n=41), 16 (n=34), 20 (n=23), and 24 hpa (n=35). (E) Two lines show the percentages of sibling and mutant NMs from (D) with at least one new HC at 12-24 hpa. (F) A diagram shows *atoh1a* regulation during pre-ablation, early-phase, and late-phase HC regeneration. Error bars in (B), (D), and (E) show SEMs, and the statistical significance between genotypes or timepoints was assessed with one-sided Mann-Whitney U tests for (B) and (D) and chi-squared tests for (E) (solid lines, p<0.05; **, p<0.01; ***, p<0.001).

Because *ybx1* is the central hub in the early GRN, we wondered whether mutants’ small but consistent deficits in HC number indicate early delays in regeneration. To test this hypothesis, we obtained *Tg(atoh1a:dTomato)* zebrafish, which we bred with heterozygous *ybx1* mutants to yield equal numbers of wild-type and heterozygous mutant offspring that expressed dTomato in newly regenerated HCs (Figure 2C). In heterozygous mutants, the average number of new HCs per NM remained around one between 12-16 hpa, whereas new HCs in sibling NMs increased to around two over the same period (Figure 2D). The proportion of sibling NMs with at least one new HC also led mutants by 4 h, reaching around 80%by 16 hpa *versus* 20 hpa in mutants (Figure 2E). These results indicate that *ybx1* lies upstream of *atoh1a* during the first 20 hrs of regeneration and can promote about 20%faster regeneration in wild-type siblings.

## Discussion

Our results indicate that *ybx1* promotes the rapid regeneration of HCs in damaged NMs and suggest a mechanism for the early transcriptional control of *atoh1a*. In mice, *Atoh1* transcription involves promoter derepression by Hes/Hey TFs—”activator insufficiency”—and 3’ enhancer-mediated autoregulation (the “Atoh switch”)^37–39^. However, initially surpassing the threshold for self-regulation requires the activity of unknown TFs at the promoter^37^. We suggest that in zebrafish *ybx1* is one such TF that initially upregulates *atoh1a* expression through interactions with the cRRPE (Figure 2F). This activity follows promoter derepression by Hes/Hey TFs (*e*.*g*., *her4*.*1* and *her4*.*2*.*1*) and anticipates the maintenance of its expression by itself and other TFs.

Even though the role of *ybx1* in upregulating *atoh1a* expression is likely restricted to cyprinids, these results strongly support our approach to supervised GRN inference with DELAY. Transfer learning can identify important patterns in noisy representations of dynamic processes such as pseudotime trajectories^28^. Our results here and elsewhere have demonstrated that DELAY is particularly effective at identifying gene-regulatory hubs whose dynamic expression either propagates downstream to genes such as *atoh1a* or arises from complex upstream regulation such as *ESRRB*^29^. Future studies can use DELAY to elucidate the causal mechanisms underlying temporal regulation of gene expression across diverse biological systems.

## Materials and methods

### Single-cell RNA-sequencing data and pseudotime inference

We used droplet-based single-cell RNA-sequencing data from neomycin-treated NMs at 0, 0.5, 1, 3, 5, and 10 hrs after HC ablation and employed the same quality-control and pre-processing steps in Seurat (version 5.0.1) as in the original study^16,40^. We then identified in the tSNE the clusters that correspond to the central SCs, HC progenitors, young HCs, and mature HCs. Prior to pseudotime inference with Slingshot (version 2.8.0), updated tSNE embeddings were computed for each trajectory^41^.

### Transfer learning and gene-regulatory inference with DELAY

Log-normalized RNA counts were used to train DELAY (version 0.2.0) and infer the HC- and SC-specific GRNs during early regeneration^28^. We trained DELAY’s RNA-specific model jointly on ground-truth interactions for both the HC and SC trajectories with targets of *sox2, sox21a, six1a*, and *six1b*^26^. The complete GRNs during early regeneration were then inferred independently for each trajectory.

### *De novo* discovery of Ybx1’s DNA-binding motif

To identify enriched motifs in *ybx1*’s predicted targets, we gathered 100 bp fragments of targets’ and controls’ 100 kb enhancer sequences for analysis (UCSC zebrafish genome GRCz11; STREME version 5.5.5)^42^. We then used FIMO (version 5.5.5) to determine the relative enrichment of Ybx1’s DNA-binding motif across predicted targets’ promoters *versus* controls^43^.

### Zebrafish husbandry, strains, and genotyping

Experiments were performed on zebrafish larvae three to five days post-fertilization. Eggs were collected and maintained at 28.5 °C in egg water containing 1 mg/L methylene blue. Heterozygous *ybx1*^*sa34489*^ zebrafish larvae were obtained from the Zebrafish International Resource Center. The following primers were used for PCR and Sanger sequencing of *ybx1* mutant offspring: 5’-GTGGAGAGATGTGACAGAATATCG-3’ (forward); 5’-CATAACTGAAATAAACCCTGGAGCG-3’ (reverse). We also used *Tg(atoh1a:dTomato)* larvae to visualize newly regenerated HCs by fluorescence microscopy.

### Ablation of HCs in posterior lateral-line NMs

To ablate all first primordium-derived HCs, we treated 3 dpf larvae twice for 1 hr each with 300 μM neomycin sulfate (Sigma-Aldrich, St. Louis, USA) with 6 hrs of recovery between treatments. We performed the second treatment to kill all non-regenerating HCs that survived or matured after the first treatment. We used 3 dpf larvae because they are less susceptible to systemic neomycin toxicity than older fish though their HCs are more susceptible to chemical ablation.

### Fixation, staining, and fluorescence imaging

To label HCs, we fixed 3-5 dpf zebrafish trunks with 4% formaldehyde in phosphate-buffered saline (PBS) solution with 0.1 % Tween-20 (0.1 % PBST) for either 1 hr at room temperature or overnight at 4 °C. For inbred *ybx1* mutant offspring, we washed trunks for 30 m in fresh 0.1 % PBST, followed by a 1 hr incubation at room temperature in 0.05 % PBST with DAPI (1:200) and Alexa Fluor Plus 405 phalloidin (1:40; Invitrogen, Eugene, USA). For *Tg(atoh1a:dTomato)* larvae, we washed trunks in 0.5 % PBST and incubated them for 1 hr at room temperature in 0.05 % PBST blocking solution with 1 % BSA and again for 2 hr in fresh blocking solution with DAPI (1:200), phalloidin (1:40) and goat anti-tdTomato DyLight550-conjugated antibodies (1:50; MyBioSource, San Diego, USA). We washed trunks once more for 20-30 m in 1X PBS before mounting and imaging at 100X on an Olympus IX83 inverted confocal microscope with a microlens-based super-resolution imaging system (VT-iSIM, VisiTech International).

## Acknowledgements

We would like to acknowledge Gaurav Shrestha, Nicolas Velez, Emily Atlas, Agnik Dasgupta, and other members of the Laboratory of Sensory Neuroscience for helpful discussions, as well as Samantha Campbell for expert zebrafish husbandry. We also thank Tatjana Piotrowski for kindly providing *Tg(atoh1a:dTomato)* larvae.

## Additional information

## Author contributions

Caleb C Reagor, Conceptualization, Methodology, Investigation, Formal analysis, Visualization, Software, Writing—original draft preparation, Writing—review & editing; A J Hudspeth, Supervision, Writing—review & editing.

## Ethics

Experiments were performed in accordance with the standards of Rockefeller University’s Institutional Animal Care and Use Committee.

## Data availability

All scripts and analyzed data for this study and pre-processed files for Gene Expression Omnibus dataset GSE196211 are available at https://github.com/calebclayreagor/ybx1-HC-regeneration. Raw fluorescent microscopy images will be available as a Dryad dataset upon publication.

## References

1. Corwin, J. T. Postembryonic production and aging of inner ear hair cells in sharks. J. Comp. Neurol. 201, 541–553 (1981).

2. Cruz, I. A. et al. Robust regeneration of adult zebrafish lateral line hair cells reflects continued precursor pool maintenance. Dev. Biol. 402, 229–238 (2015).

3. Jones, J. & Corwin, J. Replacement of lateral line sensory organs during tail regeneration in salamanders: identification of progenitor cells and analysis of leukocyte activity. J. Neurosci. 13, 1022–1034 (1993).

4. Metcalfe, W. K. Sensory neuron growth cones comigrate with posterior lateral line primordial cells in zebrafish. J. Comp. Neurol. 238, 218–224 (1985).

5. Gompel, N. et al. Pattern formation in the lateral line of zebrafish. Mech. Dev. 105, 69–77 (2001).

6. Nogare, D. D. et al. In toto imaging of the migrating Zebrafish lateral line primordium at single cell resolution. Dev. Biol. 422, 14–23 (2017).

7. Chitnis, A. B., Dalle Nogare, D. & Matsuda, M. Building the posterior lateral line system in zebrafish. Dev. Neurobiol. 72, 234–255 (2012).

8. Kindt, K. S., Finch, G. & Nicolson, T. Kinocilia Mediate Mechanosensitivity in Developing Zebrafish Hair Cells. Dev. Cell 23, 329–341 (2012).

9. Balak, K., Corwin, J. & Jones, J. Regenerated hair cells can originate from supporting cell progeny: evidence from phototoxicity and laser ablation experiments in the lateral line system. J. Neurosci. 10, 2502–2512 (1990).

10. Itoh, M. & Chitnis, A. B. Expression of proneural and neurogenic genes in the zebrafish lateral line primordium correlates with selection of hair cell fate in neuromasts. Mech. Dev. 102, 263–266 (2001).

11. Lush, M. E. et al. scRNA-Seq reveals distinct stem cell populations that drive hair cell regeneration after loss of Fgf and Notch signaling. eLife 8, e44431 (2019).

12. Montgomery, J. C., Baker, C. F. & Carton, A. G. The lateral line can mediate rheotaxis in fish. Nature 389, 960–963 (1997).

13. McHenry, M. J., Feitl, K. E., Strother, J. A. & Van Trump, W. J. Larval zebrafish rapidly sense the water flow of a predator’s strike. Biol. Lett. 5, 477–479 (2009).

14. Smith, M. E. & Monroe, J. D. Causes and Consequences of Sensory Hair Cell Damage and Recovery in Fishes. in Fish Hearing and Bioacoustics (ed. Sisneros, J. A.) vol. 877 393–417 (Springer International Publishing, Cham, 2016).

15. Harris, J. A. et al. Neomycin-Induced Hair Cell Death and Rapid Regeneration in the Lateral Line of Zebrafish (Danio rerio). JARO - J. Assoc. Res. Otolaryngol. 4, 219–234 (2003).

16. Baek, S. et al. Single-cell transcriptome analysis reveals three sequential phases of gene expression during zebrafish sensory hair cell regeneration. Dev. Cell 57, 799-819.e6 (2022).

17. Carrillo, S. A. et al. Macrophage Recruitment Contributes to Regeneration of Mechanosensory Hair Cells in the Zebrafish Lateral Line. J. Cell. Biochem. 117, 1880–1889 (2016).

18. Denans, N. et al. An anti-inflammatory activation sequence governs macrophage transcriptional dynamics during tissue injury in zebrafish. Nat. Commun. 13, 5356 (2022).

19. Romero-Carvajal, A. et al. Regeneration of Sensory Hair Cells Requires Localized Interactions between the Notch and Wnt Pathways. Dev. Cell 34, 267–282 (2015).

20. Thomas, E. D. & Raible, D. W. Distinct progenitor populations mediate regeneration in the zebrafish lateral line. eLife 8, e43736 (2019).

21. Jiang, L., Romero-Carvajal, A., Haug, J. S., Seidel, C. W. & Piotrowski, T. Gene-expression analysis of hair cell regeneration in the zebrafish lateral line. Proc. Natl. Acad. Sci. 111, (2014).

22. Bermingham, N. A. et al. Math1 : An Essential Gene for the Generation of Inner Ear Hair Cells. Science 284, 1837–1841 (1999).

23. Sarrazin, A. F. et al. Proneural gene requirement for hair cell differentiation in the zebrafish lateral line. Dev. Biol. 295, 534–545 (2006).

24. Cai, T., Seymour, M. L., Zhang, H., Pereira, F. A. & Groves, A. K. Conditional Deletion of Atoh1 Reveals Distinct Critical Periods for Survival and Function of Hair Cells in the Organ of Corti. J. Neurosci. 33, 10110–10122 (2013).

25. Chonko, K. T. et al. Atoh1 directs hair cell differentiation and survival in the late embryonic mouse inner ear. Dev. Biol. 381, 401–410 (2013).

26. Jimenez, E. et al. A regulatory network of Sox and Six transcription factors initiate a cell fate transformation during hearing regeneration in adult zebrafish. Cell Genomics 2, 100170 (2022).

27. Iyer, A. A. & Groves, A. K. Transcription Factor Reprogramming in the Inner Ear: Turning on Cell Fate Switches to Regenerate Sensory Hair Cells. Front. Cell. Neurosci. 15, 660748 (2021).

28. Reagor, C. C., Velez-Angel, N. & Hudspeth, A. J. Depicting pseudotime-lagged causality across single-cell trajectories for accurate gene-regulatory inference. PNAS Nexus 2, pgad113 (2023).

29. Mannens, C. C. A. et al. Chromatin accessibility during human first-trimester neurodevelopment. Nature (2024) doi:10.1038/s41586-024-07234-1.

30. Zasedateleva, O. A. et al. Specificity of Mammalian Y-box Binding Protein p50 in Interaction with ss and ds DNA Analyzed with Generic Oligonucleotide Microchip. J. Mol. Biol. 324, 73–87 (2002).

31. Sun, J., Yan, L., Shen, W. & Meng, A. Maternal Ybx1 safeguards zebrafish oocyte maturation and maternal-to-zygotic transition by repressing global translation. Development dev.166587 (2018) doi:10.1242/dev.166587.

32. Guarino, A. M. et al. YB-1 recruitment to stress granules in zebrafish cells reveals a differential adaptive response to stress. Sci. Rep. 9, 9059 (2019).

33. Lasham, A., Lindridge, E., Rudert, F., Onrust, R. & Watson, J. Regulation of the human fas promoter by YB-1, Pura and AP-1 transcription factors. (2000).

34. Lyabin, D. N., Eliseeva, I. A. & Ovchinnikov, L. P. YB-1 protein: functions and regulation. WIREs RNA 5, 95–110 (2014).

35. Zaucker, A. et al. Translational co-regulation of a ligand and inhibitor by a conserved RNA element. Nucleic Acids Res. 46, 104–119 (2018).

36. Weingarten-Gabbay, S. et al. Systematic discovery of cap-independent translation sequences in human and viral genomes. Science 351, aad4939 (2016).

37. Abdolazimi, Y., Stojanova, Z. & Segil, N. Selection of cell fate in the organ of Corti involves the integration of Hes/Hey signaling at the Atoh1 promoter. Development 143, 841–850 (2016).

38. Helms, A. W., Abney, A. L., Ben-Arie, N., Zoghbi, H. Y. & Johnson, J. E. Autoregulation and multiple enhancers control Math1 expression in the developing nervous system. Development 127, 1185–1196 (2000).

39. Zine, A. et al. Hes1 and Hes5 Activities Are Required for the Normal Development of the Hair Cells in the Mammalian Inner Ear. J. Neurosci. 21, 4712–4720 (2001).

40. Hao, Y. et al. Dictionary learning for integrative, multimodal and scalable single-cell analysis. Nat. Biotechnol. 42, 293–304 (2024).

41. Street, K. et al. Slingshot: cell lineage and pseudotime inference for single-cell transcriptomics. BMC Genomics 19, 477 (2018).

42. Bailey, T. L. STREME: accurate and versatile sequence motif discovery. Bioinformatics 37, 2834–2840 (2021).

43. Grant, C. E., Bailey, T. L. & Noble, W. S. FIMO: scanning for occurrences of a given motif. Bioinformatics 27, 1017–1018 (2011).

